# An intronic copy number variation in *Syntaxin 17* determines speed of greying and melanoma incidence in Grey horses

**DOI:** 10.1101/2023.11.05.565619

**Authors:** Carl-Johan Rubin, McKaela Hodge, Rakan Naboulsi, Madeleine Beckman, Rebecca R. Bellone, Angelica Kallenberg, Stephanie J’Usrey, Hajime Ohmura, Kazuhiro Seki, Aoi Ohnuma, Brian W. Davis, Teruaki Tozaki, Gabriella Lindgren, Leif Andersson

## Abstract

The Greying with age phenotype involves loss of hair pigmentation whereas skin pigmentation is not reduced and a predisposition to melanoma. The causal mutation was initially reported as a duplication of a 4.6 kb intronic sequence in *Syntaxin 17*. The speed of greying varies considerably among Grey horses. Here we demonstrate the presence of two different *Grey* alleles, *G2* carrying two tandem copies of the duplicated sequence and *G3* carrying three. The latter is by far the most common allele, probably due to strong selection for the striking white phenotype. Our results reveal a remarkable dosage effect where the *G3* allele is associated with fast greying and high incidence of melanoma whereas *G2* is associated with slow greying and low incidence of melanoma. Epigenetic analysis, based on nanopore sequencing of genomic DNA, reveals a drastic reduction in DNA methylation in part of the duplicated sequence harboring MITF binding sites. The copy number expansion transforms a weak enhancer to a strong melanocyte-specific enhancer that underlies hair greying (*G2* and *G3*) and a drastically elevated risk of melanoma (*G3* only).

## Introduction

Greying with age in horses shows dominant inheritance and is one of the most iconic mutant phenotypes in animals^1^. These horses are born fully pigmented and usually start greying during their first year of life and most eventually become completely white. The beauty of these white horses has had a major impact on human culture, white horses occur abundantly in art, sagas, and fiction, and have most certainly been an inspiration for the myths regarding Pegasus and the Unicorn. An important reason for the popularity of this phenotype is that the causal mutation only affects hair pigmentation while skin and eye pigmentation are unaffected. Additionally, there are no noticeable negative effects on vision or hearing as is the case for many other pigmentation mutations in vertebrates with pleiotropic effects. For instance, mutations in the microphthalmia-associated transcription factor gene (*MITF*) are associated with deafness in dogs^2^, Waardenburg syndrome type 2 in humans^3^, and reduced eye size and early-onset deafness in mice^4^. Grey horses are believed to be as fit as any other horse, with the exception of the melanoma prevalence described below, and are as fast as other horses as illustrated by the fact that they successfully compete in horse races with horses of other colors. The mutation is widespread and is particularly common in some breeds, for instance Lipizzaner and Arabian horses. One negative side effect, though, is that in grey horses there is a high incidence of melanomas that occur in glabrous skin with increasing incidence by age^1^ (**Extended Data Fig. 1**). The causal mutation for Grey was originally identified as a 4.6 kb duplication in intron 6 of *Syntaxin 17* (*STX17*)^1^. However, a recent study^5^, based on digital PCR, indicated that the *Grey* haplotype carries three copies of the duplicated fragment. Horses homozygous for the *Grey* mutation (*G/G*) are reported to have significantly higher melanoma incidence than heterozygotes (*G/g*)^1^. The grey melanoma is usually benign, but can grow to sizes that affect quality of life and can lead to metastasis with fatal outcomes^6-8^.

The *Grey* mutation causes upregulated expression of both *STX17* and the neighboring gene *NR4A3*, encoding an orphan nuclear receptor^1^. Previous studies using both transfection experiments in a pigment cell line and transgenic zebrafish demonstrated that the duplicated region contains a melanocyte-specific enhancer^9^. The enhancer is evolutionarily conserved and contains two binding sites for microphthalmia transcription factor (MITF), a master regulator of gene expression in pigment cells^4^. MITF knock-down silenced pigment cell-specific expression from a reporter construct with the horse copy number variation in transgenic zebrafish^9^. It is still an open question whether upregulation of *STX17* or *NR4A3* or their combined effect is critical for the phenotypic effects of the *Grey* mutation. No other study has revealed a direct link between these genes and pigment biology but both have links to cancer. The STX17 protein has a key role in autophagy^10^, a pathway currently explored target for melanoma therapy in humans^11^. The function of NR4A3 as a regulator of transcription is only partially understood, but data indicate that this orphan nuclear receptor has a role in maintaining cellular homeostasis and in pathophysiology including tumor development^12^.

It is well known that the speed of greying varies considerably among Grey horses. One important factor is the *Grey* genotype as *Grey* homozygotes grey faster than *Grey* heterozygotes^1^. However, the variation in the speed of greying is pronounced in certain breeds such as Connemara ponies. In this breed two distinct types of greying are noted, fast greying and slow greying (**Fig. 1**). Fast greying horses are usually completely white at about 10 years of age whereas the slow greying horses never become completely white but show a beautiful dappled grey color at older ages. The aim of the present study was to explore the genetic basis for this striking phenotypic difference. Here we report that this variation is controlled by the *Grey* locus itself rather than by other genes. The causal difference is the copy number of the duplicated sequence, fast greying horses carry at least one allele with three copies of the 4.6 kb sequence while slow greying horses carry an allele with only two copies. Thus, the *Grey* locus harbors an allelic series with at least three alleles: *G1* - one copy, wild type; *G2* – two copies, slow greying; *G3* – three copies, fast greying (**Fig. 2**).

**Fig. 1.**
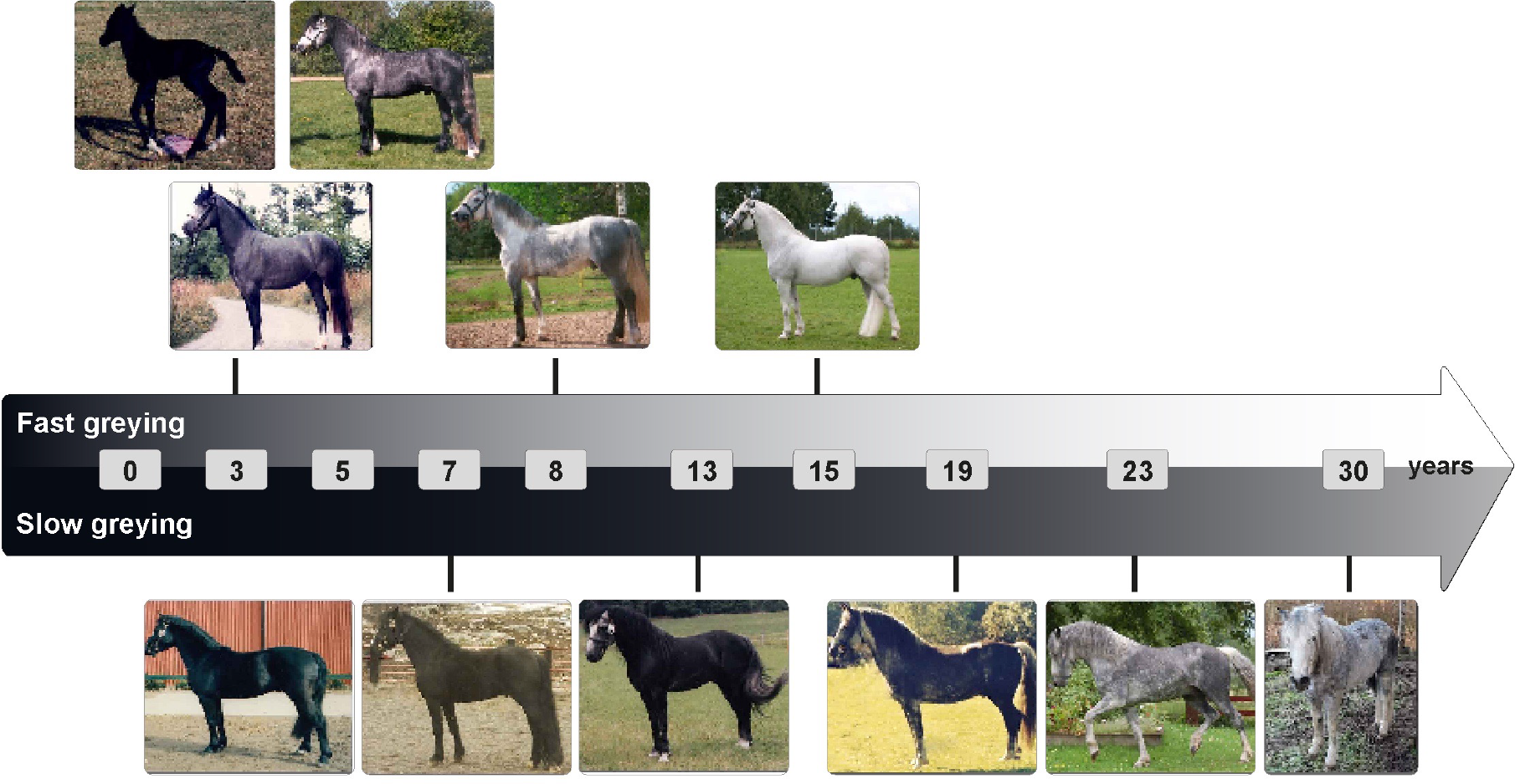
Illustration of fast and slow greying in Connemara Ponies. Two individual horses, one fast greying (upper timeline) and one slow greying (lower timeline), were used for the illustration. Photo, from left to right: Fast Grey 1,2 Madeleine Beckman, 3 Maria Johansson, 4,5 Beckman Archive. Slow Grey 1-4 Maria Johansson, 5-6 Marlen Näslin.

**Fig. 2.**
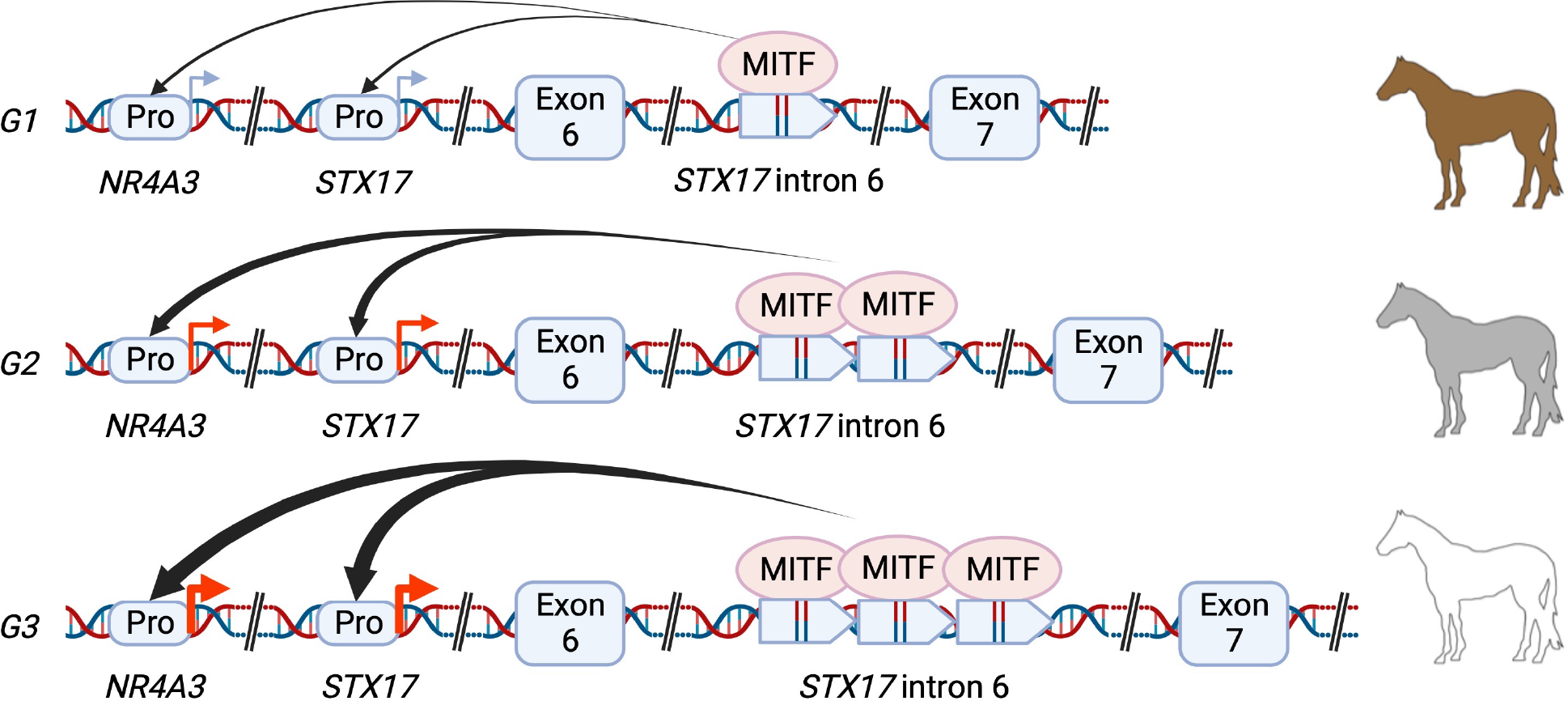
Schematic illustration of the nature and significance of the *Grey* mutation. The causal mutation for the Grey phenotype is a copy number variation of a 4.6 kb sequence located in *STX17* intron 6. The duplicated sequence harbors two binding sites for MITF and the presence of copy number expansion upregulates the expression of both *NR4A3* and *STX17* in cis^5^. *G1* is the wild-type allele with a single copy of the 4.6 kb sequence, G2 carries two copies and causes slow greying, and *G3* carries three copies and causes fast greying^5^.

## Results

### Speed of greying maps to the *Grey* locus

We identified a family of Connemara ponies apparently segregating for fast and slow greying (**Extended Data Fig. 2**). We took advantage of this pedigree in an attempt to identify the genetic basis for the difference in speed of greying. The key individual was a fast greying sire Hagens D’Arcy (stud book number RC 101) in generation 5 whose mother was fast greying and whose father greyed slowly. Among the 16 progeny from matings to non-grey (*G1/G1*) dams, 10 were classified as slow greying while 6 were fast greying. This suggests that the speed of greying in this family may show a Mendelian inheritance and the observed proportion of the two types did not differ from an expected 1:1 ratio if the sire is heterozygous for alleles associated with fast and slow greying (*Χ*^*2*^=1.0, d.f. =1; *P*>0.05). We also had access to a small family of Thoroughbred horses from Japan comprising two horses classified as fast greying and three as slow greying (**Extended Data Fig. 3**).

The putative locus controlling speed of greying may be due to sequence variation at the *STX17/Grey* locus itself, a linked locus, or an unlinked locus. To distinguish these possibilities, we carried out whole genome sequencing of a total of 13 slow greying and 7 fast greying horses, including the Connemara pony sire Hagens D’Arcy. A genome-wide association analysis using individual SNPs revealed a single consistent signal of association on chromosome 25 harboring the *STX17/Grey* locus (**Fig. 3a**). Further characterization of the signal of association revealed that it was centered around the *STX17*/*Grey* locus (**Fig. 3b**).

**Fig. 3.**
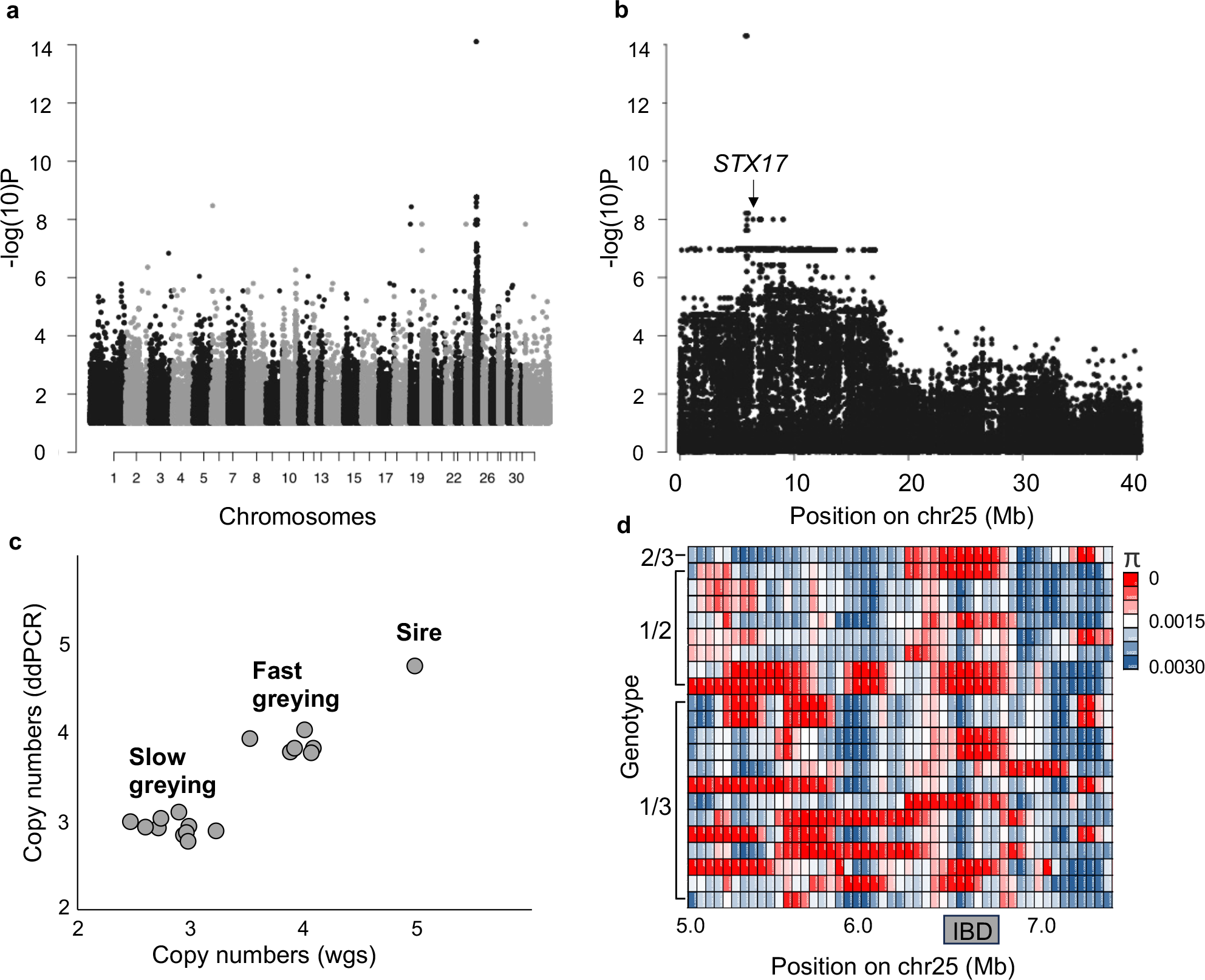
Whole genome sequencing reveals copy number difference at the *STX17/Grey* locus between fast and slow greying horses. (**a**) GWAS contrasting fast and slow greying horses belonging to the Connemara Pony breed and Japanese Thoroughbred horses. (**b**) GWAS zoom in on chromosome 25. The location of *STX17* is marked with an arrow. (**c**) Estimated copy number of the *STX17* copy number variation in slow greying and fast greying progeny of the sire Hagens D’Arcy (*G/G*) mated to non-grey (*g/g*) mares. (d) Nucleotide diversity in windows of 100 kb in the region flanking the *STX17* gene. Deduced *Grey* genotypes are indicated to the left. The sire Hagen’s D’Arcy is row 1 and he shows homozygosity for the 350 kb IBD region shared by *Grey* haplotypes.

We next hypothesized that the causal difference between fast and slow greying could be caused by a copy number difference of the 4.6 kb duplicated sequence in *STX17*. We determined the copy number both by sequence coverage in the whole genome sequencing data and by droplet digital PCR of the duplicated sequence first using samples from the Connemara pony family (**Supplementary Table 1**). This analysis showed that the fast greying sire Hagens D’Arcy carried 5 copies of the duplicated sequence (deduced to be 3+2 copies on the two chromosomes), the fast greying progeny carried 4 copies (3+1 because their dams are wild-type and transmit an allele with a single copy), and the slow greying progeny carried 3 copies (2+1) **Fig. 3c**). The analysis of the sequence coverage obtained for the five Japanese Thoroughbred horses also confirmed the same pattern. The perfect match between copy number of the duplicated sequence and the fast/slow greying phenotype including both Connemara ponies from Sweden and Thoroughbreds from Japan is highly significant (Fisher’s exact test; *P*<0.0001).

The great majority of, if not all, Grey horses previously studied and determined to be fast greying appear to carry *Grey* haplotypes that share a 350 kb region that is identical-by-descent (IBD)^1,13^. This implies that the initial mutation can be traced back to a single mutational event that happened subsequent to horse domestication. An important question is therefore whether the two-copy *Grey* haplotype shares the same IBD region as the three-copy haplotype. The sequence data from all Grey horses included in this study were consistent with the presence of this IBD region in haplotypes both with 2 and 3 copies of the duplicated sequence strongly suggesting that the two alleles have a common origin and that copy number variation has evolved after the initial duplication (**Fig. 3d**). This is illustrated by the fact that Hagens D’Arcy, which is heterozygous for the two haplotypes, shows essentially no sequence variation at the 350 kb IBD region. Interestingly also three *G1/G2* and three *G1/G3* heterozygotes show a similar pattern indicating that a very similar haplotype occurs among wild-type haplotypes carrying a single copy of the 4.6 kb sequence. An interesting possibility is that these may have originated by back mutations in which 2-copy or 3-copy haplotypes have lost the duplication or triplication by unequal crossing-over.

Based on these results we propose a new nomenclature for the *Grey* locus: *G1* - one copy, wild type, previously reported as N by most genetic testing laboratories; *G2* – two copies, slow greying; *G3* – three copies, fast greying (**Fig. 2**).

### Targeted long read sequencing supports the causal nature of the copy number variation

Nanopore (ONT) Cas9-targeted sequencing (nCATS) makes it possible to sequence chromosome fragments of tens of kb without amplification and thereby establishing structure and methylation status of individual haplotypes, including the sequence orientation of duplicated fragments^14^. We used this approach to characterize the two grey haplotypes of Hagens D’Arcy from the Connemara pony family; the genotype of this sire is *G2*/*G3*. ONT sequencing of DNA fragments spanning the duplicated region including about 2 kb of flanking sequences on each side demonstrated that the sire is heterozygous *G2/G3* based on the detection of two dominant fragments sizes: a 13 kb sequence comprising 2 copies of the duplicated sequence and a 17 kb sequence comprising 3 copies (**Fig. 4a**). After generating separate consensus sequences of reads corresponding to the G2 and G3 alleles, alignment of these sequences to the horse reference genome assembly (EquCab3) shows that the duplicated copies both in *G2* and in *G3* are located in tandem and are oriented head-to-tail (**Fig. 4b**). ONT sequencing makes it possible to also analyze the DNA methylation pattern of the sequence. Interestingly, these data revealed that the evolutionary conserved region harboring MITF binding sites^9^ shows a clear reduction in DNA methylation using genomic DNA isolated from blood (**Fig. 4c**), a result supporting our previous interpretation that this region acts as an enhancer and that its copy number variation is causal for the phenotypic effects^9^.

**Fig. 4.**
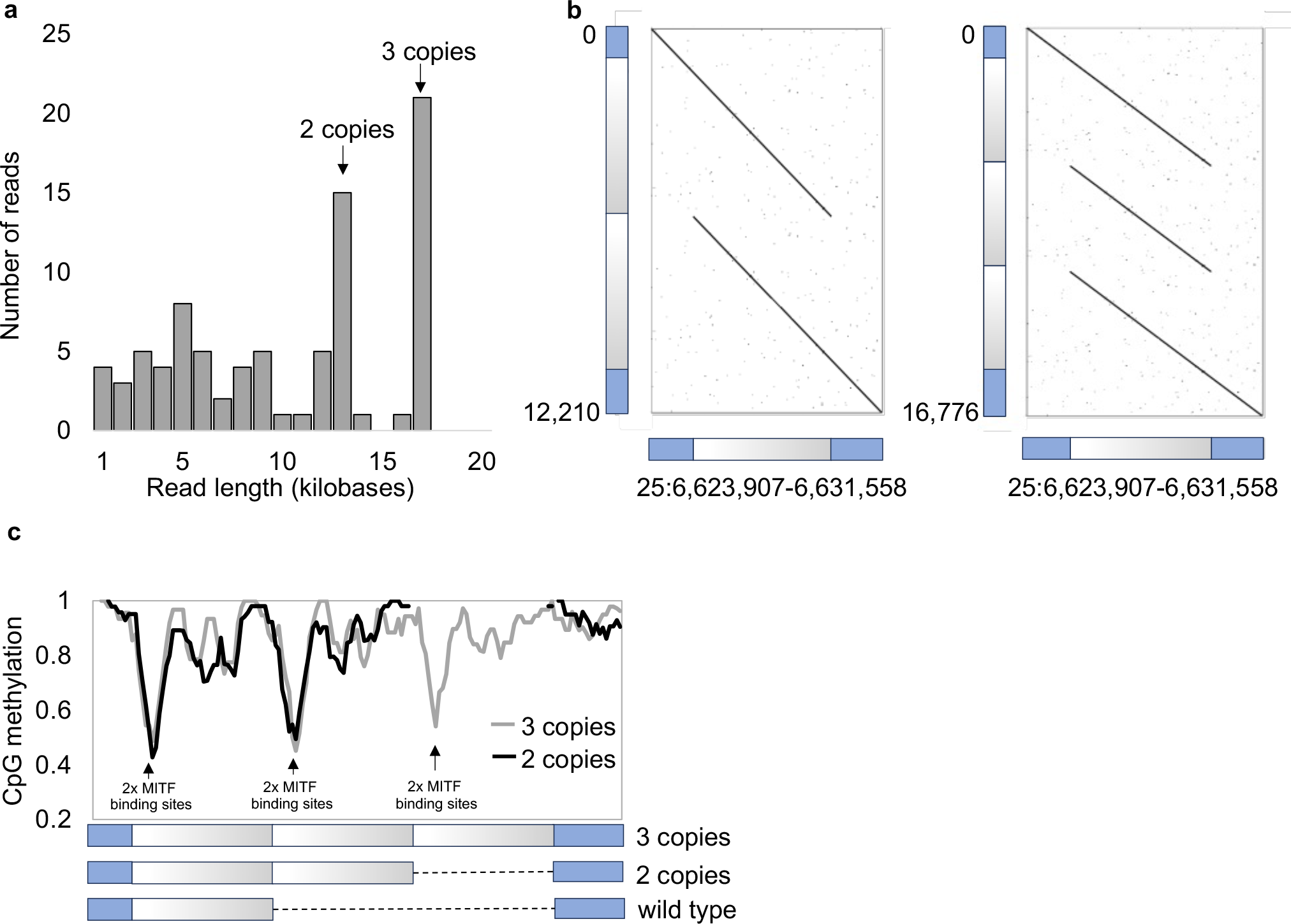
Copy number variation and DNA methylation patterns revealed by sequence capture and Nanopore sequencing. (**a**) Molecular size distribution of the *STX17* region from a *G2/G3* heterozygote (Hagens D’Arcy) inferred from a Cas9 capture Nanopore sequencing experiment. (**b**) The duplicated sequences are present in tandem both on the **G2** and **G3** alleles. (**c**) DNA methylation pattern associated with the *G2* (red) and *G3* (blue) alleles. The drastic reduction of DNA methylation at precisely the region harboring MITF binding sites are high-lighted.

Pedigree analysis identified an individual as a putative *G2/G2* homozygote. The genotype was confirmed by ONT adaptive sampling sequencing. The sequence analysis of the entire chromosome 25 revealed that this horse showed runs of homozygosity over a 16 Mb region including the *Grey* locus (chr25:2-18Mb) suggesting that both *G2* haplotypes trace back to a relatively recent common ancestor. The phenotype of this single horses was classified at the age of 11 years as intermediate between a slow greying (*G1/G2*) and a fast greying horse (*G1/G3*) (**Extended Data Fig. 4**) and it had no visible melanoma.

### Genotype distribution across breeds

There is an interest from horse owners and breeders to determine the genotype at the *Grey* locus in horses. During the last six years the Veterinary Genetics Laboratory (UC Davis) has utilized a droplet digital PCR assay to test for *Grey* genotype. Including data from those samples whose owner or registry consented to be included in research allowed us to evaluate 1,400 horses representing 78 breeds/populations. These data enabled the determination of the grey copy number distribution across breeds. The majority of samples tested had four copies (n=831). It is not possible to know from this assay alone if these animals are homozygous for the *G2* allele or are *G1/G3* heterozygotes. However, despite non-random sampling, in those breeds where we did not identify a single *G1/G2* heterozygote it is extremely unlikely that all individuals with four copies are homozygous *G2/G2*. Under this assumption, the *G3* allele was identified in 62 different populations (**Supplementary Table 2**). Whereas the *G2* allele was only definitively detected in 8 of these 62 populations (3 and/or 5 copies detected in that population screened, Table 1). Under the assumption that the genotype distribution at this locus does not show an extreme deviation from Hardy-Weinberg equilibrium, the *G2* allele was always much rarer than *G3* in the population where evidence of both alleles occurred, because horses with 4 copies were always more abundant than those with three copies in the ddPCR screens. We did not find evidence for the presence of an allele with 4 or more copies which would be evident if a horse has 7 or more copies in total. Our data indicate that if such alleles exist, they must be very rare.

**Table 1.** Copy number variation for the 4.6 duplicated sequence in *STX17* intron 6 in which the *G2* and *G3* alleles were detected. Data based on genotyping services at the UC Davis Veterinary Genetics Laboratory.

| Breed | Copy number (deduced genotype) |  |  |  |  | n | Allele frequency<sup>1</sup> |  |
| --- | --- | --- | --- | --- | --- | --- | --- | --- |
|  | 2 | 3 | 4 | 5 | 6 |  | *G2* | *G3* |
|  | (*G1/G1*) | (*G1/G2*) | (*G1/G3* or<br>*G2/G2*) | (*G2/G3*) | (*G3/G3*) |  |  |  |
| Andalusian | 25 | 8 | 40 | 1 | 9 | 83 | 0.05 | 0.36 |
| Connemara Pony | 23 | 6 | 49 | 3 | 9 | 90 | 0.05 | 0.39 |
| Miniature Horse | 25 | 1 | 15 | 0 | 0 | 41 | 0.01 | 0.18 |
| Mangalarga Marchador | 0 | 1 | 8 | 0 | 0 | 9 | 0.06 | 0.44 |
| Mustang | 3 | 2 | 8 | 0 | 0 | 13 | 0.08 | 0.31 |
| Quarter Horse | 141 | 0 | 284 | 1 | 13 | 439 | 0.00 | 0.35 |
| Tennessee Walking Horse | 16 | 2 | 23 | 0 | 4 | 45 | 0.02 | 0.34 |
| Welsh Pony | 7 | 1 | 25 | 0 | 3 | 36 | 0.01 | 0.43 |

<sup>1</sup>Allele frequency calculated under the assumption that no *G2/G2* homozygote is present among these samples due to the low expected frequency determined by the number of *G1/G2* and/or *G2/G3*. These are biased estimates of allele frequencies in these breeds but give an indication of the relative frequency of the *G2* and *G3* alleles.

### The slow greying phenotype is not associated with a high incidence of melanoma

Our observation that the *G2* allele has a much milder effect on the greying process than the *G3* allele begs the question whether it is also associated with a lower incidence of melanoma. We identified 25 late grey (*G1/G2*) Connemara horses, 15 years of age or older (range 15-35 years) and searched for the presence of the classical grey melanomas present on glabrous skin (lips, eye lids, under the tail) but detected none among these horses (**Supplementary Table 3**). Previous studies have documented that from 50 - 80% of all Grey horses of this age carry melanoma^6-8^. This dramatic difference in incidence 80% vs. 0% in 25 slow-greying horses is highly significant (*Χ*^*2*^=25.0, d.f. =1; *P*=6x10^-7^).

## Discussion

This study provides conclusive evidence for the presence of two variant alleles at the *Grey* locus in horses in addition to the *G1* wild-type allele. The most common allele, *G3*, carries three tandem copies of the 4.6 kb sequence and is associated with fast greying that eventually results in a white coat color and the allele is associated with a high incidence of melanoma. In contrast, the *G2* allele carries a tandem duplication and *G1/G2* heterozygotes grey slowly, never become pure white and have a significantly lower incidence of melanoma compared with *G1/G3* heterozygotes, possibly as low as *G1/G1* homozygotes. The fact that *G2* and *G3* share a 350 kb IBD region strongly suggests that the two alleles originate from the same ancestral mutation. It is unknown if the ancestral allele carried two or three copies of the 4.6 kb sequence, but it is likely that these alleles can be generated from each other by unequal crossing-over or slippage during DNA replication. As evidenced in our across breed analysis, *G3* is much more common than *G2* and this is likely caused by strong selection for the splendid white phenotype associated with this allele. Our results now solve the mystery why some grey horses never become pure white. Although this study explains a considerable portion of the heterogeneity in speed of greying, it is likely that additional genetic variation elsewhere in the genome contributes to variation in the speed of greying and these additional loci may also contribute to melanoma risk. It has been shown that in addition to the large effect of the *STX17* mutation, polygenic inheritance contributes to the speed of greying and incidence of melanoma in Lipizzaner horses^15^.

There is a remarkably strong dose sensitivity at the *Grey* locus. The *G1/G1* genotype shows no greying and low incidence of melanoma, *G1/G2* shows slow greying and low incidence of melanoma, *G1/G3* shows fast greying and high incidence of melanoma, and *G3/G3* homozygotes grey very fast and show a very high incidence of melanoma^1^. There are too few observations to make firm predictions of the phenotype of *G2/G3* heterozygotes and *G2/G2* homozygotes. However, it is likely that *G2/G3* horses are intermediate compared with the *G1/G3* and *G3/G3* genotypes with regard to hair greying and melanoma incidence. The single *G2/G2* individual confirmed by sequence capture and Nanopore sequencing showed a grey phenotype and no melanoma at the age of 11 years whereas a *G1/G3* heterozygote is expected to have developed white color and is likely to have some melanoma at the same age. This suggests that having a triplication on one chromosome gives a stronger transcriptional activation than having a duplication on both chromosomes. The present study has important practical implications for horse breeding because it suggests that it is possible to have a horse showing the beauty of the grey coat color (**Fig. 1**) most likely with no elevated risk to develop melanoma. It also illustrates the need for additional testing methodologies to determine zygosity of this tandem repeat in those horses with four copies (*G2/G2* or *G1/G3*). Current testing methodologies, which do not routinely use long read sequencing for many reasons, are therefore not yet able to resolve this issue.

Our results provide strong support for the causality of the copy number variation on the Grey phenotype, because of the remarkable correlation between the copy number of the 4.6 kb sequence and phenotype. It also supports the interpretation that the triplication underlies both loss of hair pigmentation through its effect on hair follicle melanocytes and melanoma incidence through its effect on dermal and/or epidermal melanocytes. The higher copy number enhances the effect on both speed of greying and melanoma risk. Furthermore, *G2* alleles from two different breeds (Connemara Pony and Thoroughbreds) share a 350 kb IBD region with *G3* alleles from many different breeds making the copy number of the duplicated sequence the distinctive difference between alleles. The 4.6 kb sequence showing copy number variation contains two binding sites for MITF, a master regulator for transcription in pigment cells^4^. A previous study based on both cellular transfection experiments and transgenic zebrafish indicated that this copy number expansion transforms a weak enhancer to a strong melanocyte-specific enhancer^9^. The drastic drop in DNA methylation precisely at the region where the MITF binding sites are located (Fig. 4c) despite the fact that we used genomic DNA from blood cells for nanopore sequencing is consistent with the notion that the 4.6 kb sequence contains an element affecting gene regulation. It is likely that the *STX17* copy number variation is a direct driver of melanoma development because we noted further copy number expansion in melanoma cells in horses, up to nine copies in tumor tissue^13^. A classical feature of tandem duplications is expansions and contractions of the copy number in the germ line due to non-allelic homologous recombination^16^. Thus, since we observe haplotypes with both two and three copies it is expected that haplotypes with four or more germ-line copies may occur. Genotyping 1,400 horses representing 78 populations has not revealed further expansion of this tandem repeat beyond three copies in the germ line. This observation together with our finding of low DNA methylation of the enhancer region also in blood cells and the strong dosage effect suggest that four or more tandem copies of the duplicated sequence may be deleterious and thus exist at low frequency, if at all.

## Methods

### Pedigree data

The Connemara pedigree from Sweden consisted of the sire Hagens D’Arcy and 16 of his offspring from matings to non-grey mares. Five fast greying horses (three Arabians, one Swedish warmblood (SWB), and one Dutch warmblood (KWPN) and five non-Grey horses (three Swedish warmblood, one Dutch warmblood (KWPN), and one Icelandic horse) from other horse breeds were used as controls (see **Supplementary Table 1**). Blood sample collection was approved by the ethics committee for animal experiments in Uppsala, Sweden (numebr: 5.8.18-15453/2017 and 5.8.18-01654/2020). Genomic DNA from all horses were extracted on the Qiasymphony instrument with the Qiasymphony DSP DNA mini or midi kit (Qiagen, Hilden, Germany).

### Grey phenotyping

All Grey horses, fast or slow greying, are born with a primary fully pigmented coat, but the onset and progress of greying differ markedly between slow and fast greying horses. A fast greying foal will normally display a few grey hairs on the eyelids during the first week after birth (**Extended Data Fig. 5a**). During the first month more grey hairs appear as grey “circles” around the eyes (**Extended Data Fig. 5b**), grey eyelashes and grey hairs on the tail root develop too. In contrast, a slow greying foal has no visible grey hairs on the eyelids at birth and does not develop grey “eye circles” or any grey hairs elsewhere during its first month of life. If a slow greying foal has face markings (like a star or blaze) the white parts are not well-defined and the white hairs will spread out in time (**Extended Data Fig. 5c**). The first signs of greying in a slow greying horse will normally not appear until 5 to 7 years of age, with grey hairs often expanding from white head markings (**Extended Data Fig. 5d**). Grey eyelashes can appear when the head is still dark colored with no grey hairs on the eyelids. Grey hairs may appear partly on the neck. The next phase of greying will include head, neck, distal parts of the leg and the tail. When a slow greying horse has reached an age of about 15 years the coat color may turn into a dappled steel or bluish grey shade. Although there is individual variation in the speed of greying also among slow greying horses, they will never become a visually white horse (**Fig. 1**); **Extended Data Fig. 5e, f and g** show the slow greying process of the same horse at different ages, but without head markings. The breeder of the slow greying *G2/G2* homozygote from the Connemara Pony in our study stated that the horse had developed the slow greying color like a normal slow Grey Connemara Pony, but that it had turned grey earlier than expected, but not at all as early as a fast greying heterozygote (*G1/G3*). In contrast to heterozygous slow greying horses (*G1*/*G2*), it started with greying of the head much like a fast greying horse. As a three-year-old, it began to have grey hair on its head and it lacked white eyelashes. At 8 years old, it had a light head but the body was still dark. Only when it was 11 years old the grey color developed on the body and the characteristic dappled colored pattern began to appear on the front of the horse (**Extended Data Fig. 4**). The head is still lighter but overall, the horse differs markedly from a fast greying horse of the same age.

### Whole genome sequencing

A Covaris E220 system was used to fragment 1 μg of genomic DNA to a target insert size of 350-400bp. Sequencing libraries of individual DNA samples were prepared using the Illumina TruSeq DNA PCR-free LP kit (Illumina, San Diego, California, United States) in combination with the IDT for Illumina TruSeq DNA UD indexes kit (Illumina). Preparation of the libraries was performed according to the manufacturer’s protocol. A TapeStation with the D1000 ScreenTape (Agilent Technologies, Santa Clara, California, USA) was used to assess the quality of the libraries. qPCR was performed using the Library quantification kit for Illumina (KAPA Biosystems, Basel, Switzerland) on a CFX384 Touch instrument (Bio-Rad, Hercules, California, USA) to quantify the adapter-ligated fragments prior to cluster generation and sequencing. Indexed samples were sequenced on a NovaSeq S4 flow cell and the resulting demultiplexed sequences were aligned to the horse genome (EquCab3) using bwa-mem2^17^. Read duplicates were marked using MarkDuplicates^18^ and resulting bam-files were used for variant calling by UnifiedGenotyper^18^. Depth of coverage was determined by means of the GATK module DepthOfCoverage and the resulting per-base coverage files were used to calculate mean depths of coverage across samples for windows of varying sizes. In order to estimate DNA copy numbers in windows, depths of coverage in individual windows were normalized to the genome average depth.

### Genome Wide Association (GWAS) analysis

Genotype data in vcf format was loaded into the Python API for GWAS analysis vcf2gwas^19^, which uses algorithms from GEMMA^20^ for association tests. GWAS was performed using a univariate linear mixed model with phenotypes of 13 fast greying individuals encoded as “5” and phenotypes of 8 slow greying individuals encoded as “10”. Resulting P values were imported into R^21^ and were plotted along the genome using the package qqman^22^. Genotypes of three SNPs on chromosome 25 were perfectly associated with the trait and their P-values for association were more than a hundred orders of magnitude lower than those of associated SNPs with a non-perfect association. For the purpose of easy interpretation of Manhattan plots, -log(10P) values or these three positions were changed to a value of 14 before plotting in qqman. The three perfectly associated positions were chr25:5747798, chr25:5743983 and chr25:5868699.

### Nanopore Cas9-targeted sequencing

High molecular weight DNA was isolated from blood of one *G2/G3* individual. We used a Cas9 sequence capture protocol (Ligation sequencing gDNA - Cas9 enrichment; SQK-CS9109) to construct a sequencing library where DNA molecules spanning the 4.6 kb duplication/triplication were much more likely than other fragments to ligate with the sequencing adapter. Sequencing was performed on a R9.4.1 flow cell using a MinION instrument. The Cas9 Sequencing Kit (SQK-CS9109) was purchased from Oxford Nanopore Technologies. We designed four guideRNAs (**Supplementary Table 3**) and purchased these as S. pyogenes Cas9 Alt-R™ crRNAs (Integrated DNA Technologies). Alt-R™ S. pyogenes Cas9 tracrRNA (Integrated DNA Technologies) and Alt-R S. pyogenes HiFi Cas9 nuclease V3 (Integrated DNA Technologies) were used to cut DNA. Base calling was performed using Guppy Version 6.5.7 (Oxford Nanopore Technologies) using the super accuracy setting. Reads corresponding to the G2 and G3 alleles, carrying two or three copies of the duplication, were separately binned and consensus sequences were determined for each of these. Individual G2 and G3 reads were then aligned to their matching consensus sequence using minimap2^23^ and the resulting bam files and read fast5 files were used to call CpG methylation in nanopolish^24^.

### Nanopore sequencing by adaptive sampling

High molecular weight DNA was isolated from blood of one *G2/G2* individual. This DNA was fragmented to 20 kb using a g-TUBE (Covaris) and molecules below approximately 10 kb in size were then depleted using the SRE XS kit (Pacific Biosciences of California). For sequencing library construction, the kit SQK-LSK114 was used and 40 fmol of the finished library was loaded onto a PromethION flow cell (Oxford Nanopore Technologies) for sequencing. The run was started from within MinKNOW on a P2 instrument with adaptive sampling enabled to achieve high coverage on chr25. Adaptive sampling used the horse reference genome fasta file (EquCab3) and a bed-file specifying chromosomes 1-24 and chromosomes 26-27 (88% of the genome) as off-target coordinates.

### Digital PCR and copy number analysis

Horses from the Swedish Connemara pony pedigree and controls were genotyped using two TaqMan assays designed to specifically target the *STX17* copy number variation (ECA25: 6625371-6625493 (NC_009168.3), Bio-Rad Assay dCNS718018816; and ECA25: 6629062-6629184 (NC_009168.3), Bio-Rad Assay dCNS933010968) and a third control assay to target a neighboring region free of known CNVs (ECA25: 6624137-6624259 (NC_009168.3), Bio-Rad Assay dCNS851288843. A fourth assay targeting *myostatin (MSTN)*^25^ was used as a reference in the ddPCR experiment. The assays were designed using EquCab3 as a reference genome. The ddPCR experiment was performed using the Bio-Rad QX200 Droplet Reader platform. Briefly, droplets were generated using the Bio-Rad Automated Droplet Generator instrument before placing the ddPCR mix in a thermal cycler where the PCR reaction was performed. Finally, the results were generated using Bio-Rad QX200 Droplet Reader platform. The ddPCR mix contained 11 μL of 2X ddPCR supermix for probes, 1.1 μL of target 20X TaqMan assays, 1.1 μL of reference 20X TaqMan assays, 1 μL of 20 ng/μL genomic DNA, 2 μL fast digest EcoRI, and water up to 22 μL. The mix was loaded into a droplet generator cartridge. Droplets were generated following the manufacturer’s protocol. The PCR plate containing the droplets was foil sealed before the PCR reaction. The PCR setup was 95°C for 10 min, 40 cycles of 30 s at 94°C and 60 s at 60°C. Then finally, 10 min at 98°C. The PCR plate was transferred to the droplet reader for analysis. Cluster classification was manually adjusted in non-clear events. Additionally copy number was evaluated for 1400 horses across 78 breeds based on genotyping services provided by the UC Davis Veterinary Genetics Laboratory.

### Incidence of melanoma in slow greying horses

All 16 horses that were sampled in the study (10 slow gray (*G1/G2*) and 6 fast gray (*G1/G3*) offspring descending from the stallion Hagens D’Arcy with non-grey mothers) were examined clinically by a veterinarian regarding the presence of melanoma. Two of the fast greying offspring (11 and 13 years old, respectively) were found to be affected by melanoma on the underside of the tail, while the rest of the offspring were negative. As the sampled horses were relatively young (half were between 0-7 years old) and the total number low, it was not possible to draw any conclusions regarding the melanoma incidence for fast grays and slow grays respectively in the Connemara pony based on this material.

To estimate the incidence of melanoma in slow greying Connemara ponies, we did a study of 25 slow greying horses between 15 and 35 years old (**Supplementary Table 3**). Four of the horses were included in the mapping study as offspring of Hagens D’Arcy and were examined by a veterinarian for melanoma at the time of blood sampling. The examination of the other 21 horses was carried out as an interview where a veterinarian asked neutral questions to owners or breeders of the horses if they had observed visible melanomas in the horses from the age of 15 onwards. Those who answered the questions were experienced breeders or horse keepers who were judged to have good knowledge of the symptoms of melanoma in horses. None of the respondents stated that they had observed melanoma in the horses in question.

## Author contributions

LA and GL conceived the study. C-JR was responsible for nCATS, nanopore adaptive sampling and bioinformatic analyses. MH, BWD and C-JR designed and MH, BWD evaluated the nCATS experiment. RN performed ddPCR analysis and prepared sequencing libraries for the Connemara pony family. MB collected family material and phenotype data from Connemara ponies. RB, AK, and SJU were responsible for *Grey* genotyping across breeds. HO, KS, AO, and TT were responsible for collection of the Japanese Thoroughbred family. LA wrote the manuscript with input from other authors. All authors approved the manuscript before submission.

## Data availability statement

The sequence data generated in this study have been submitted to NCBI (http://www.ncbi.nlm.nih.gov/PRJNA1035120).

## Code availability statement

The analyses of data have been carried out with publicly available software and all are cited in the Methods section. Code associated with bioinformatic analyses will be available at: https://github.com/LeifAnderssonLab/

## Competing interest statement

R.R. Bellone, A. Kallenberg, and S. J’Usrey are affiliated with the UC Davis Veterinary Genetics Laboratory, which provides genetic diagnostic tests in horses and other species. The other authors declare no competing interest.

## Acknowledgements

We thank all horse owners, in particular the owners of Hagens D’Arcy and his offspring that have allowed us to sample their horses and consented to research. We also thank Dr. Robert Grahn, Thea Ward, Shayne Hughes, Elizabeth Esdaile, and Tytti Vanhala for their technical assistance. The project was financially supported by Vetenskapsrådet (2017-02907; to LA), Knut and Alice Wallenberg Foundation (KAW 2016.0361; to LA). The National Genomics Infrastructure (NGI)/Uppsala Genome Center provided service in massive parallel sequencing and the computational infrastructure was provided by the Swedish National Infrastructure for Computing (SNIC) at UPPMAX partially funded by the Swedish Research Council (2018-05973).

## Author information

## Supplementary material

**Extended Data Fig. 1.**
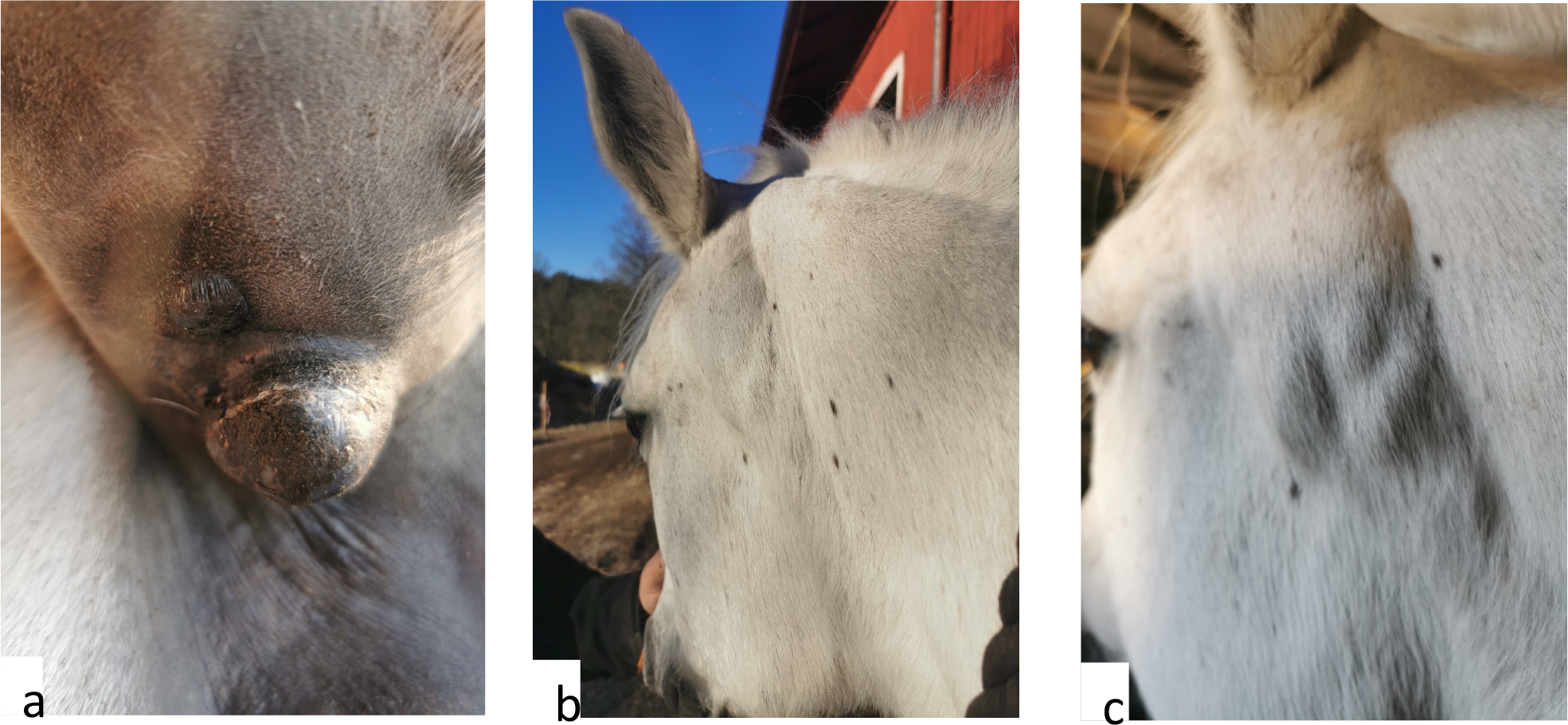
Melanoma in grey Connemara Ponies. The most common locations of visual melanoma in Grey horses are under the tail and in the jaw angel (parotis area). (**a**) Prominent melanoma under the tail of a 12 years old Connemara mare. The father is Hagens D’Arcy and the mother is a random fast greying horse. (**b**) Melanoma in the jaw angle accumulated around the lymphatic tissue (most likely metastasis) of a 15 years old Connemara mare. The father is Hagens D’Arcy and the mother is a random fast greying mare. (**c**) A zoomed in view of Extended Data Fig. 1b where the lymphatic tissue-associated melanoma visually bulges under the skin. The two melanoma displaying mares were not part of the mapping pedigree since their mothers were fast greying horses. Photo: Elisabeth Ljungstorp.

**Extended Data Fig. 2.**
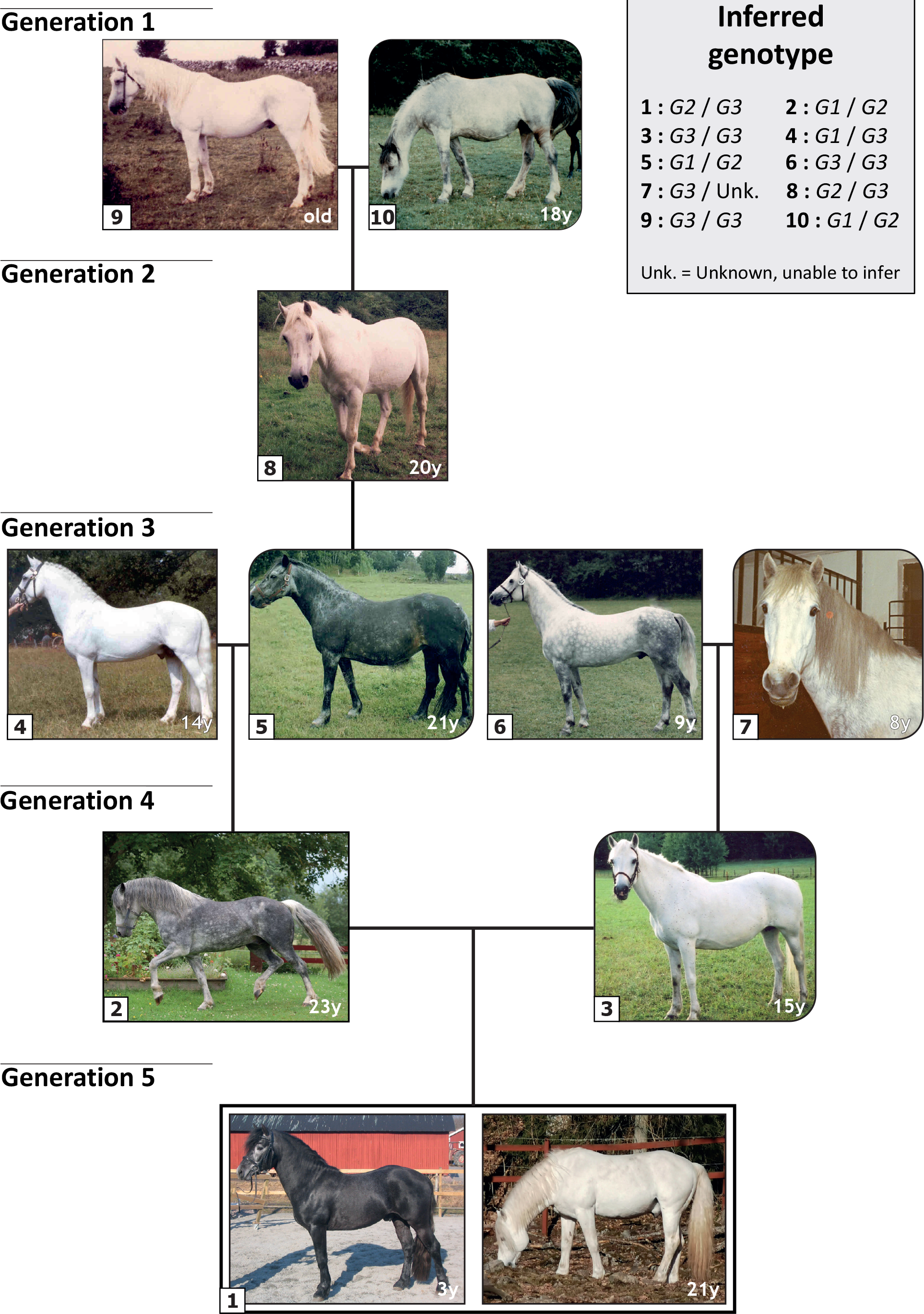
Five generation pedigree of Connemara Ponies segregating for slow and fast greying. The sire in generation 5 **(Hagens D’Arcy)** was postulated to be heterozygous for slow/fast greying because his parents were fast greying (mother) and slow greying (father) and in matings with non-grey dams he had progeny that were either classified as slow or fast greying. The inferred genotypes of all horses in the pedigree are based on information listed in **Supplementary Table 5**. Photo: 1 Elisabeth Ljungstorp, 2 Marlen Näslin, 3 Madeleine Beckman, 4 Anneli Dahlskog, 5-8 Madeleine Beckman, 9 Beckman Archive, 10 Madeleine Beckman.

**Extended Data Fig. 3.**
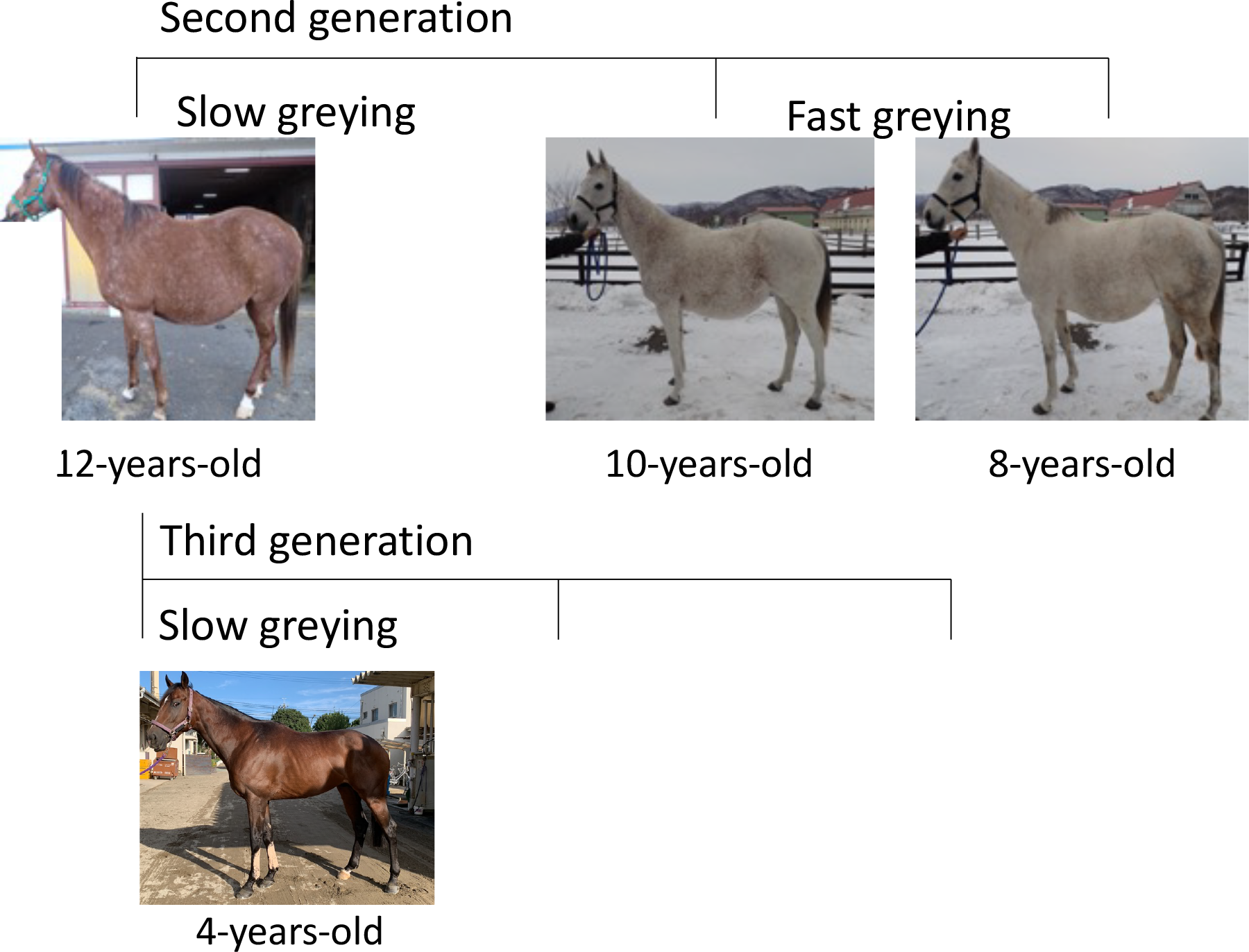
Three generation pedigree of Japanese Thoroughbred horses segregating for fast and slow greying. The dam in the first generation was fast greying according to the Japanese Thoroughbred Stud Book. She produced one slow greying and two fast greying progenies in matings with non-grey sires (second generation). Three slow greying horses from the third generation were offspring of the slow greying dam (generation two) in matings with non-grey sires. Photos (two 3-years-old horses in third generation) were provided by the Japan Racing Association.

**Extended Data Fig. 4.**
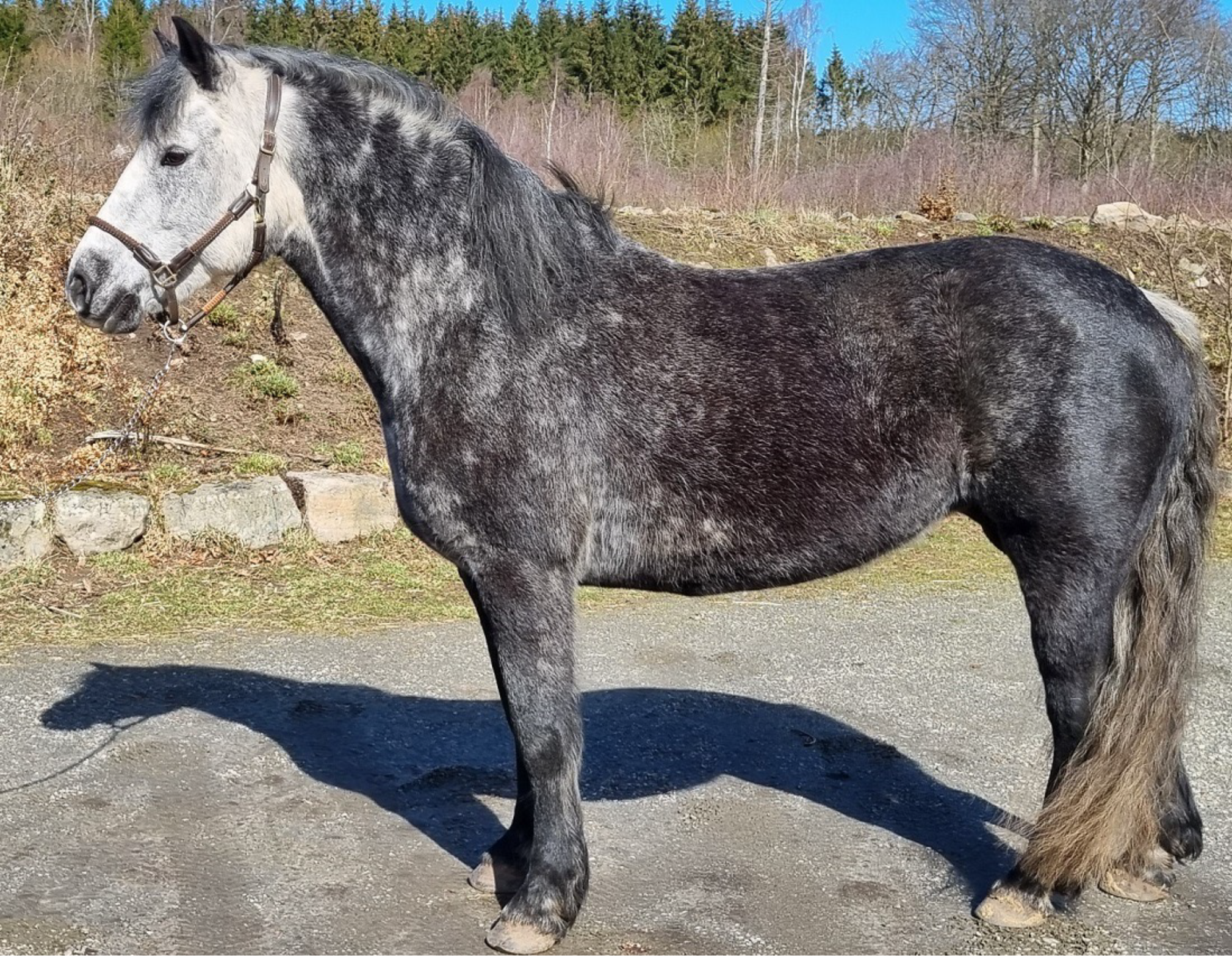
Phenotype of a *G2/G2* homozygote from the Connemara pony breed at the age of 11 years. Photo: Madeleine Beckman.

**Extended Data Fig. 5.**
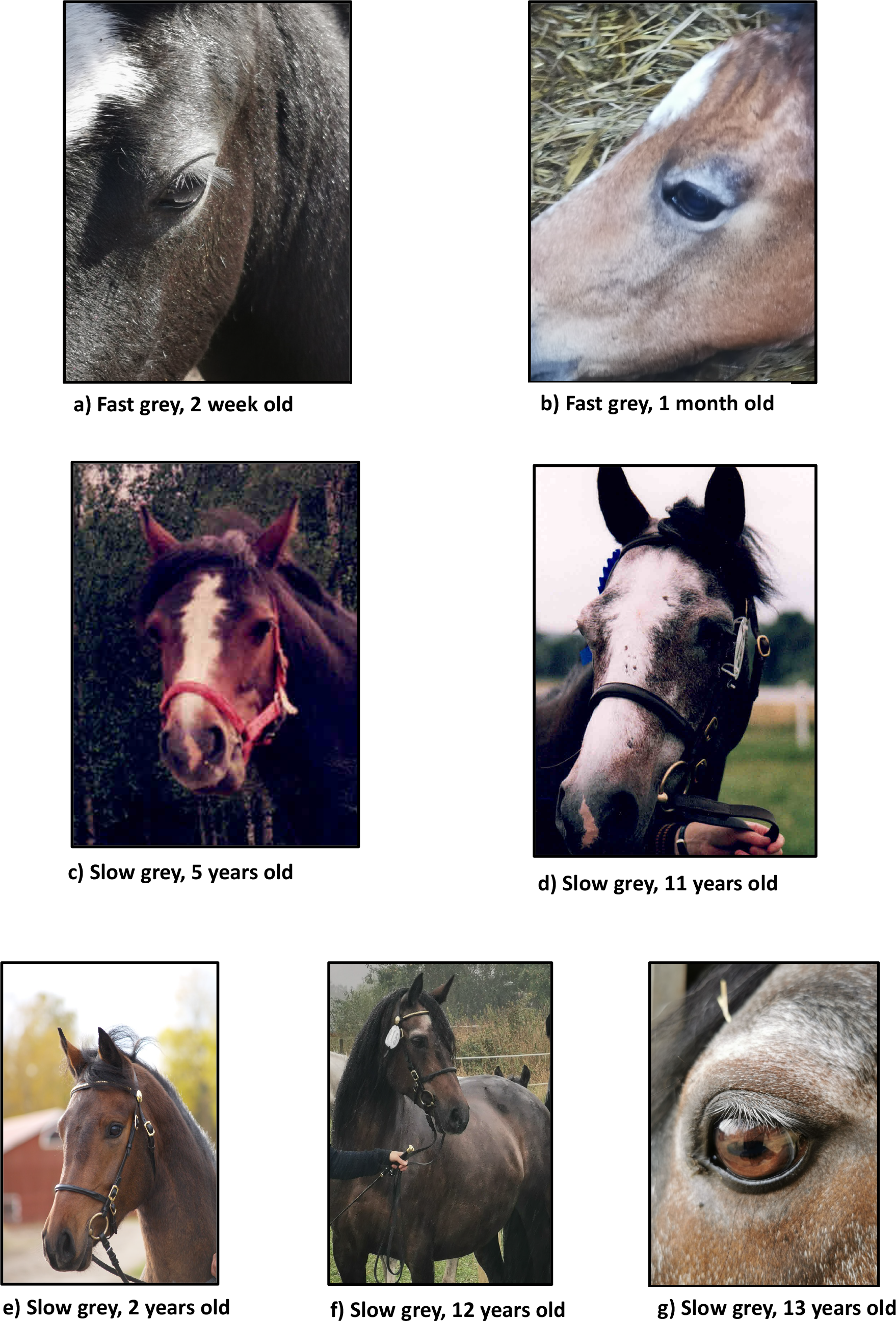
Comparison of the coat color phenotype of fast and slow greying Connemara ponies from younger age and onwards. **(a)** Fast greying foal 2 weeks old. Grey hairs visible on the eyelids. (**b**) Fast greying foal about 1 month old. Circles of grey hairs around the eyes. (**c, d**) Slow greying mare at 5 and 11 years, showing a white blaze with shattering edges, slowly spreading out. (**e, f, g**) Slow greying mare at 2, 12, and 13 years. Pure bay with no visible grey hairs at 2 years of age. Eyelashes turning grey between 12 and 13 years of age. Photo: Elisabeth Ljungstorp (a, b, e, f, g) and Madeleine Beckman (c, d).

**Supplementary Table 1. Results of a ddPCR experiment** performed with two TaqMan assays targeting the *STX17* copy number variation (CNV) (Assays A and B) and a control assay (Assay C) targeting an *STX17* region that is free of known CNV. Copy number for these samples were also validated and confirmed using the UC Davis VGL assay.

**Supplementary Table 2. Copy number variation for the 4.6 sequence in *STX17* intron 6 in 1400 horses across 78 populations. Data based on genotyping services provided at the UC Davis Veterinary Genetics Laboratory. (Excel file)**

**Supplementary Table 3. Age distribution of 25 slow greying (*G1/G2*) Connemara ponies examined for the incidence of melanoma. None was diagnosed with melanoma**.

**Supplementary Table 4. Guide RNAs used for Cas9 sequence capture of the *STX17* duplicated region**.

**Supplementary Table 5. Inferred *Grey* genotypes for the horses included in Extended Data Fig. 2**.

