## Supplementary Tables for "An intronic copy number variation in *Syntaxin 17* determines speed of greying and melanoma incidence in Grey horses"

**Supplementary Table 1**. Results of a ddPCR experiment performed with two TaqMan assays targeting the *STX17* copy number variation (CNV) (Assays A and B) and a control assay (Assay C) targeting an *STX17* region that is free of known CNV.

| | | | Copy number | |
|---|---|---|---|---|
| **Horses** | **Breed** | **Phenotype** | **CNV assay** | **Control assay** |
| Hagens D'Arcy | Connemara | Fast greying | 4.75 | 2.02 |
| Offspring 1 | Connemara | Fast greying | 3.77 | 1.91 |
| Offspring 2 | Connemara | Slow greying | 2.92 | 1.96 |
| Offspring 3 | Connemara | Slow greying | 3.03 | 1.96 |
| Offspring 4 | Connemara | Fast greying | 3.82 | 1.99 |
| Offspring 5 | Connemara | Slow greying | 2.99 | 1.89 |
| Offspring 6 | Connemara | Fast greying | 3.78 | 1.9 |
| Offspring 7 | Connemara | Slow greying | 2.84 | 2.00 |
| Offspring 8 | Connemara | Slow greying | 2.93 | 1.97 |
| Offspring 9 | Connemara | Slow greying | 2.87 | 1.95 |
| Offspring 10 | Connemara X Welsh | Slow greying | 2.94 | 1.92 |
| Offspring 11 | Connemara | Slow greying | 2.77 | 1.98 |
| Offspring 12 | Connemara | Fast greying | 3.82 | 2.05 |
| Offspring 13 | Connemara X SWB | Slow greying | 2.89 | 2.03 |
| Offspring 14 | Connemara | Slow greying | 3.10 | 1.98 |
| Offspring 15 | Connemara X SWB | Fast greying | 3.93 | 2.02 |
| Offspring 16 | Connemara | Fast greying | 4.03 | 1.96 |
| Control 1 | KWPN | Fast greying | 3.74 | 1.94 |
| Control 2 | Arabian | Fast greying | 5.77 | 1.90 |
| Control 3 | SWB | Fast greying | 4.39 | 2.03 |
| Control 4 | Arabian | Fast greying | 6.24 | 1.99 |
| Control 5 | Arabian | Fast greying | 3.96 | 1.97 |
| Control 6 | SWB | Non-grey | 1.88 | 1.97 |
| Control 7 | Icelandic horse | Non-grey | 2.05 | 2.05 |
| Control 8 | SWB | Non-grey | 1.94 | 1.96 |
| Control 9 | SWB | Non-grey | 2.00 | 1.95 |
| Control 10 | KWPN | Non-grey | 2.06 | 1.96 |

KWPN=Dutch warmblood; SWB=Swedish warmblood.

**Supplementary Table 2. Copy number variation for the 4.6 sequence in *STX17* intron 6 in 1400 horses across 78 populations. Data based on genotyping services provided at the UC Davis Veterinary Genetics Laboratory.**

| | Copy Number | | | | | Sample Size |
|---|---|---|---|---|---|---|
| **Breed** | **2** | **3** | **4** | **5** | **6** | |
| Andalusian | 25 | **8** | 40 | **1** | 9 | 83 |
| Appaloosa | 6 | | 10 | | | 16 |
| Arabian | 12 | | 87 | | 40 | 139 |
| American Sport Pony | | | 1 | | | 1 |
| Appendix | 2 | | 3 | | | 5 |
| Azteca | 2 | | 1 | | 1 | 4 |
| British Spotted Pony | 1 | | | | | 1 |
| Belgian Warmblood | 1 | | | | | 1 |
| Caballo Deportivo la Silla | 1 | | | | | 1 |
| Connemara Pony | 23 | **6** | 49 | **3** | 9 | 90 |
| Caspian Horse | | | 4 | | 2 | 6 |
| CR Iberoamericano | 1 | | 4 | | 1 | 6 |
| Costa Rican Paso Horse | 1 | | 14 | | 2 | 17 |
| Continental Warmblood | | | 1 | | | 1 |
| Canadian Warmblood | | | 1 | | | 1 |
| Dales Pony | 2 | | 3 | | | 5 |
| Deutsches Reitpony | 2 | | | | | 2 |
| Deutsches Sportpferd | | | 1 | | | 1 |
| Dutch Warmblood | 2 | | 3 | | | 5 |
| Draft Cross | 1 | | | | | 1 |
| Brabant | 3 | | | | | 3 |
| Fell Pony | | | | | 1 | 1 |
| Friesian Sporthorse | 3 | | | | | 3 |
| French Warmblood (Selle Francais) | | | 1 | | | 1 |
| Friesian Cross | 3 | | 2 | | | 5 |
| Gypsy Cob | 1 | | 2 | | | 3 |
| German Riding Pony | | | 1 | | | 1 |
| Gypsy Vanner | 3 | | 1 | | | 4 |
| Half Andalusian | | | 1 | | | 1 |
| Hanoverian | 1 | | 2 | | 1 | 4 |
| Holsteiner | 1 | | 5 | | 1 | 7 |
| Iberian Warmblood | 1 | | | | | 1 |
| Icelandic Horse | 2 | | 1 | | | 3 |
| Irish Cob | | | 1 | | | 1 |
| Irish Draught | | | 5 | | 2 | 7 |
| Irish Sport Horse | 2 | | | | | 2 |
| Knabstrupper | 2 | | | | | 2 |
| Lipizzaner | 2 | | 13 | | 21 | 36 |
| Lusitano | 11 | | 26 | | 15 | 52 |
| Moroccan Barb | | | | | 3 | 3 |
| Morgan Horse | 10 | | 16 | | 1 | 27 |
| Miniature Horse | 25 | **1** | 15 | | | 41 |
| Mangalarga Marchador | | **1** | 8 | | | 9 |
| Mustang | 3 | **2** | 8 | | | 13 |
| Mule | 1 | | | | | 1 |
| Norwegian Fjord Horse | 3 | | | | | 3 |
| Nokota | 1 | | 1 | | | 2 |
| New Forest Pony | | | 1 | | | 1 |
| National Show Horse | | | 1 | | | 1 |
| Oldenburg | 3 | | 7 | | 1 | 11 |
| Other | 4 | | 9 | | 1 | 14 |
| Pony of the Americas | | | 1 | | | 1 |
| Paso Fino | 1 | | | | | 1 |
| Pinto | | | 3 | | | 3 |
| Percheron | 4 | | 9 | | 2 | 15 |
| Paint Quarter Horse | 2 | | | | | 2 |
| Pura Raza Española | 8 | | 14 | | 2 | 24 |
| Paint Horse | 2 | | 6 | | | 8 |
| Quarter Horse | 141 | | 284 | **1** | 13 | 439 |
| Racking Horse | 1 | | | | | 1 |
| Rheinland | 1 | | 1 | | | 2 |
| Rocky Mountain Horse | 4 | | 4 | | 2 | 10 |
| Single-Footing Horse | 1 | | 2 | | | 3 |
| Shire | 3 | | 34 | | 1 | 38 |
| Shetland Pony | 4 | | 5 | | 2 | 11 |
| Spanish Heritage Horse | | | 1 | | | 1 |
| Standardbred | | | 2 | | | 2 |
| Swedish Warmblood | 1 | | 1 | | | 2 |
| Thoroughbred | 2 | | 8 | | 1 | 11 |
| Turbo Friesian | | | 1 | | | 1 |
| Trakehner | 3 | | 11 | | 3 | 17 |
| Tennessee Walking Horse | 16 | **2** | 23 | | 4 | 45 |
| Unknown | 11 | | 16 | | 3 | 30 |
| Westfalen | 1 | | 1 | | | 2 |
| Warlander | 2 | | | | | 2 |
| Welsh Pony | 7 | **1** | 25 | | 3 | 36 |
| Crossbred | 12 | | 28 | | 2 | 42 |
| Zangersheide | | | 2 | | | 2 |
| **Grand Total** | **394** | **21** | **831** | **5** | **149** | **1400** |

**Supplementary Table 3**. Age distribution of 25 slow greying (*G1/G2*) Connemara ponies examined for the incidence of melanoma. None of the horses was diagnosed with melanoma.

| Horse | Age at observation |
|---|---|
| 1 | 35 |
| 2 | 33 |
| 3 | 21 |
| 4 | 15 |
| 5 | 18 |
| 6 | 19 |
| 7 | 22 |
| 8 | 17 |
| 9 | 16 |
| 10 | 15 |
| 11 | 16 |
| 12 | 21 |
| 13 | 16 |
| 14 | 17 |
| 15 | 17 |
| 16 | 22 |
| 17 | 22 |
| 18 | 27 |
| 19 | 28 |
| 20 | 18 |
| 21 | 22 |
| 22 | 15 |
| 23 | 22 |
| 24 | 17 |
| 25 | 15 |

**Supplementary Table 4**. Guide RNAs used for Cas9 sequence capture.

| Coordinates (EquCab3) | Sequence | Strand |
|---|---|---|
| chr25:6623878-6623900 | ATCCTGGGAAACCTTAGAAG | + |
| chr25:6623747-6623769 | GCAAATAAACTCTAAACATG | + |
| chr25:6631636-6631658 | TCACAGCAGCAGTGTTGACA | - |
| chr25:6631564-6631586 | TCACAGCAGCAGTGTTGACA | - |

**Supplementary Table 5. Inferred *Grey* genotypes for the horses included in Extended Data Fig. 2.**

| Horse | Genotype | Comments |
|---|---|---|
| 1 | *G2/G3* | Only fast or slow greying foals from 25 matings to non-grey mares |
| 2 | *G1/G2* | Only slow greying or non-grey foals from 33 matings to non-grey mares |
| 3 | *G3/G3* | 10 foals, all fast greys, 8 of them by non-grey stallions |
| 4 | *G1/G3* | Fast greying individual with non-grey sire |
| 5 | *G1/G2* | Slow greying individual with 16 foals, 4 of them where non-grey |
| 6 | *G3/G3* | 87 registered foals, all fast greys |
| 7 | *G3*/Unk. | 17 foals all fast grey, 9 of them by non-grey stallions but no clear phenotype data available on 4 of them |
| 8 | *G2/G3* | Sire *G3/G3*, Dam *G2/G1*. Approximately 70 foals registered in Ireland, both fast and slow greying individuals |
| 9 | *G3/G3* | Approximately 300 foals in Ireland, very few non-greys registered (not parentage tested). 31 foals registered in Sweden, all fast greys. |
| 10 | *G1/G2* | Slow greying individual with non-grey sire |

Unk. = unknown. Not possible to infer the genotype.
